# Segzoo: a turnkey system that summarizes genome annotations

**DOI:** 10.1101/2023.10.03.559369

**Authors:** Mickaël Mendez, Yushan Liu, Marc Asenjo Ponce de León, Michael M. Hoffman

## Abstract

Segmentation and automated genome annotation (SAGA) techniques, such as Segway and ChromHMM, assign labels to every part of the genome, identifying similar patterns across multiple genomic input signals. Inferring biological meaning in these patterns remains challenging. Doing so requires a time-consuming process of manually downloading reference data, running multiple analysis methods, and interpreting many individual results.

To simplify these tasks, we developed the turnkey system Segzoo. As input, Segzoo only requires a genome annotation file in browser extensible data (BED) format. It automatically downloads the rest of the data required for comparisons. Segzoo performs analyses using these data and summarizes results in a single visualization.

**Availability and Implementation:** Source code for Python ≥3.7 on Linux freely available for download at https://github.com/hoffmangroup/segzoo under the GNU General Public License (GPL) version 2. Segzoo is also available in the Bioconda package segzoo: https://anaconda.org/bioconda/segzoo.

## 1 Introduction

Segmentation and automated genome annotation (SAGA) methods^14^, such as Segway ^10^ and ChromHMM^6^, partition the genome into segmentation made of non-overlapping segments. The methods assign a label to each segment such that the segments sharing the same label exhibit similarities in the input data. As the input data can consist of complex combinations of chromatin immunoprecipitation sequencing (ChIP-seq) data^3,11,17^ or RNA sequencing (RNA-seq) data, inferring a biological role for a label requires comparing the segmentation against multiple reference datasets. This requires a non-trivial amount of toil ^16^, including downloading reference data. It also involves orchestrating the use of these data with a variety tools, such as Segtools^4^ and BEDTools^15^, potentially through the development of custom scripts. Interpretation requires simultaneous consideration of multiple visualizations and analyses.

To help assign biological hypotheses to SAGA labels, we designed the turnkey system Segzoo. Segzoo requires only a genome segmentation file as input to generate all the visualizations. Segzoo uses Go Get Data (GGD) ^5^ to automatically download all required data for these analyses and produces an easy to interpret figure which reveals patterns of segmented regions. Here, we demonstrate the use of Segzoo to quickly identify promoter features from a Segway annotation of the H1 human embryonic stem cell line.

## 2 Methods

Segzoo summarizes genome annotation through a four-step workflow. First, it uses GGD to download reference data for the genome assembly of the segmentation. Second, Segzoo generates segmentation-centric summary statistics using Segtools and BEDTools^15^. Third, Segzoo parses the output of each tool and summarizes them into one table. Fourth, Segzoo plots the table as a compact figure to allow rapid biological hypothesis assignment to SAGA labels.

### 2.1 Dependencies

In addition to a segmentation, Segzoo needs a gene annotation and a reference genome to perform analyses. Segzoo uses GGD to automatically download the data dependencies and Conda^2^ to automatically install the software dependencies. Segzoo uses the Snakemake^12^ workflow manager to automatically perform the necessary steps. Using Snakemake also allows taking advantage of multiprocessing to accelerate the analyses.

### 2.2 Analysis tools

Segzoo uses Segtools to perform a series of analyses. It combines the results of `segtools-aggregation`, `segtools-length-distribution`, `segtools-overlap`, and `segtools-gmtk-parameters` into a single plot. Segzoo also uses BEDTools’s ^15^ `bedtools nuc` command to calculate the nucleotide frequency and G+C content for each segmentation label.

### 2.3 Input data

Here, we demonstrate the use of Segzoo by creating a Segway annotation of the H1 human embryonic stem cell line. To do this, we used ENCODE ChIP-seq data for CTCF (ENCFF332TNJ), H3K27me3 (ENCFF417VQQ), H3K4me1 (ENCFF584AVI), H3K27ac (ENCFF919FBG), and H3K4me3 (ENCFF760NUN). We trained the 10-label Segway model with the following parameters: `segway train --resolution=10 --minibatch-fraction=0.01 --num-labels=10 --num-instances=10`. We set the environment variable SEGWAY_RAND_SEED=22426492 to make the results reproducible.

We used Segzoo to generate a detailed summary figure (Figure 1) with the following single command:

**Figure 1.**
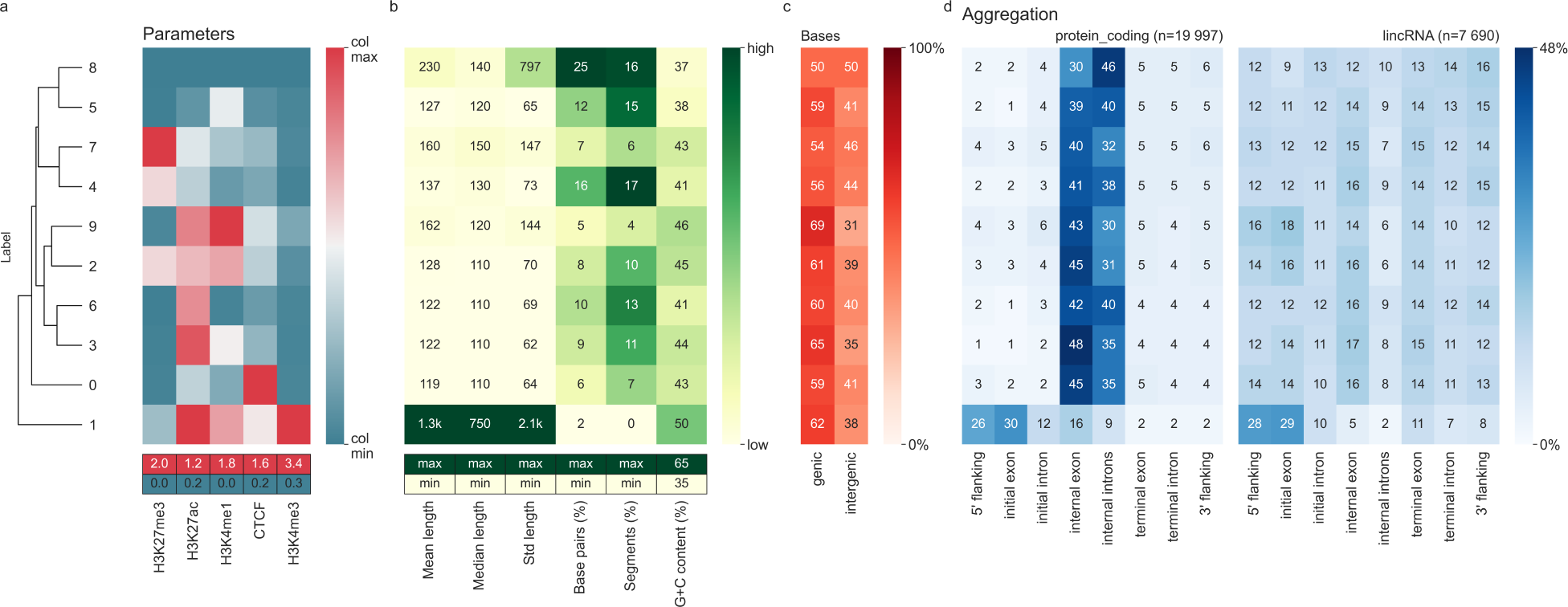
Segzoo summary of Segway H1 human embryonic stem cell line annotation. Numeric values in each heatmap represent various summary statistics for a label. **(a)** *Top:* Gaussian mean parameters from Segway trained model. Column-normalized values, with minimum and maximum values mapped to the range 0 to 100. *Left:* Dendrogram indicating hierarchical clustering on rows from weighted pair group method with arithmetic average (WPGMA) ^18^. *Bottom:* Maximum and minimum values of each column. **(b)** *Top:* Heatmap of statistics summarizing segments per label: mean segment length (“mean length”), median segment length (“median length”), standard deviation of segment length (“std length”), percentage of base pairs annotated with the label (“base pairs”), percentage of segments annotated with the label (“segments”), G+C content. The letter “k” indicates thousands. *Bottom:* Indication of maximum and minimum values of the color scale for each column from one of: column maximum value (“max”), column minimum value (“min”), other value (a number). **(c)** Fractional overlap of each label with genic and intergenic regions. **(d)** Fractional overlap between segments and components and flanking regions of GENCODE^8^ protein-coding genes (left) and long intergenic non-coding RNA (lincRNA) genes (right). Row-normalized values per table, with values summing up to 100.

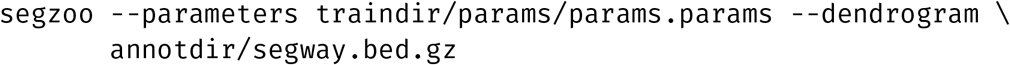

To plot the Parameters heatmap (Figure 1a), we used the `--parameter` option with Segway’s trained parameter file, `params.params`. The file contains the mean of the Gaussian distribution learned by Segway for each pair of dataset and label. The other heatmaps (Figure 1b–d) used the output of `segway annotate`, `segway.bed.gz`.

## 3 Results

The Segzoo visualization consists of an array of heatmaps, with rows representing SAGA labels and columns representing different quantitative descriptions of those labels (Figure 1). The first heatmap, “Parameters”, indicates the mean parameters of the Gaussian distributions learned for each label on each input dataset (Figure 1a). The other heatmaps summarize other aspects of the output segmentation, including summary statistics of the segments (Figure 1b), proportion of bases in genic regions (Figure 1c), and overlap between segments and components of GENCODE genes—both protein-coding genes (Figure 1d, left), and lincRNAs (Figure 1d, right). To simplify the interpretation of labels with similar patterns, Segzoo automatically performs row-wise and column-wise hierarchical clustering on the Parameters table. All the heatmaps share the same row order to simplify the interpretation of all the descriptions for a given label.

Here, Segzoo allows us to quickly see that label 1 has several promoter features (Figure 1). First, label 1 has the highest mean H3K4me3 signal (Figure 1a). This signal often indicates promoter regions ^13^. Second, label 1 has the highest G+C content, another feature of promoter regions ^1^ (Figure 1b). Label 1 has the highest mean length and median length. We don’t expect this for a promoter region. Label 1, however, has the highest length standard deviation and the fewest segments. This suggests that Segway annotated not only promoter regions, but also long genomic regions with the label 1. Third, the Overlap table shows the segments with the label 1 mostly annotate genic regions (Figure 1c). Fourth, the Aggregation table shows an enrichment of label 1 at the 5^*′*^ end of both protein coding genes and lincRNAs (Figure 1d). The four heatmaps together help generate the hypothesis that label 1 annotates promoter regions. Interestingly, we can also see easily that label 9 has promoter features such as high H3K4me3 signal, high G+C content, and frequent overlap at lincRNA 5^*′*^ ends.

## 4 Code and data availability

Segzoo supports Python ≥3.7 on Linux. Code is freely available under GNU General Public License (GPL) version 2 at https://github.com/hoffmangroup/segzoo. The Bioconda^7^ package segzoo (https://anaconda.org/bioconda/segzoo) makes it easy to install.

We have deposited in Zenodo the version of the Segzoo source which produced the results in this article (https://doi.org/10.5281/zenodo.8333102), other code and data used to produce the results (https://doi.org/10.5281/zenodo.8333120), and the results (https://doi.org/10.5281/zenodo.8333124).

## 5 Conclusion

Segzoo simplifies the process of assigning a biological hypothesis to SAGA labels. To simplify the process, it generates a compact visualization that summarizes the overlap between a SAGA annotation and multiple reference datasets. Segzoo automatically download the reference datasets via standardized recipes with GGD, which makes it easier to attain higher degrees of reproducibility. For example, this work meets the silver standard of reproducibility ^9^ .

## Acknowledgments

This work was supported by the Natural Sciences and Engineering Research Council of Canada (RGPIN-2015-03948 to M.M.H.). We also thank Carl Virtanen and Zhibin Lu at the University Health Network High-Performance Computing Centre and Bioinformatics Core for technical assistance.

